# Understanding the microbial biogeography of ancient human dentitions to guide study design and interpretation

**DOI:** 10.1101/2021.08.16.456492

**Authors:** Zandra Fagernäs, Domingo C. Salazar-García, Azucena Avilés, María Haber, Amanda Henry, Joaquín Lomba Maurandi, Andrew Ozga, Irina M Velsko, Christina Warinner

## Abstract

The oral cavity is a heterogeneous environment, varying in factors such as pH, oxygen levels, and salivary flow. These factors affect the microbial community composition and distribution of species in dental plaque, but it is not known how well these patterns are reflected in archaeological dental calculus. In most archaeological studies, a single sample of dental calculus is studied per individual and is assumed to represent the entire oral cavity. However, it is not known if this sampling strategy introduces biases into studies of the ancient oral microbiome. Here, we present the results of a shotgun metagenomic study of a dense sampling of dental calculus from four Chalcolithic individuals from the southeast Iberian peninsula (ca. 4500-5000 BP). Inter-individual differences in microbial composition are found to be much larger than intra-individual differences, indicating that a single sample can indeed represent an individual in most cases. However, there are minor spatial patterns in species distribution within the oral cavity that should be taken into account when designing a study or interpreting results. Finally, we show that plant DNA identified in the samples may be of environmental origin, showing the importance of including environmental controls or several lines of biomolecular evidence.

## Introduction

Dental calculus forms when the dental plaque biofilm mineralizes during life (Jin and Yip 2002), a periodic occurrence that encapsulates microbes, host biomolecules, food residues, and particles from the environment (Velsko and Warinner 2017). After the death of an individual, biomolecules within dental calculus can be preserved for tens of thousands of years (Fellows Yates *et al.* 2021), largely protected from environmental processes within the mineral matrix. Studies of archaeological dental calculus have rapidly increased in number in recent years, in part due to an elevated interest in the evolution of the oral microbiome and a growing understanding of the plethora of ancient biomolecules and information that can be recovered from this semi-fossilized microbial biofilm. However, there are still many unknown factors regarding the formation and preservation of archaeological dental calculus, and further method development is therefore necessary.

In carrying out comparative studies of ancient dental calculus, researchers aim to set up a sampling strategy that mitigates biases caused by intra-individual variability of the studied individuals. However, as archaeological dental calculus is often found in small quantities, especially in individuals dating far back in time, and is not always present on the same teeth across individuals, it may not always be possible to adhere to such a sampling scheme. Pre- and post-mortem tooth loss can further complicate sampling designs, as does working with calculus samples that were dislocated from the teeth during handling or storage, such that the precise tooth of origin is unknown. Due to such sampling constraints, some studies have pooled and homogenized calculus from several teeth for analysis (Warinner *et al.* 2014), which may partly mitigate spatial biases, but this approach requires the presence and collection of larger amounts of calculus, which is a finite archaeological substrate. In light of these challenges, most ancient oral microbiome studies implicitly assume that a single sample can be representative of the entire dentition, regardless of the tooth niche from which the calculus sample is obtained, and analyze only a single dental calculus deposit per individual. The oral cavity, however, is not a uniform environment, and thus microbial communities may vary across the dentition, potentially leading to bias when comparing across individuals from whom different teeth were sampled.

Differences in the microbial composition of different oral tissues, such as buccal mucosa, keratinized gingiva, saliva, and teeth, have been reported in present-day humans (Aas *et al.* 2005; Ding and Schloss 2014; Eren *et al.* 2014; Mark Welch *et al.* 2016; Proctor *et al.* 2018; Utter *et al.* 2020). Further, differences in dental plaque microbial communities have been previously reported between mandibular and maxillary teeth (Haffajee *et al.* 2009; Simon-Soro and Tomás 2013), between tooth position (e.g. anterior vs. posterior teeth) (Haffajee *et al.* 2009; Proctor *et al.* 2018), between tooth surfaces (e.g. buccal vs. lingual) (Simon-Soro and Tomás 2013; Proctor *et al.* 2018), and between supragingival and subgingival plaque (Simon-Soro and Tomás 2013; Eren *et al.* 2014). Local variations in oral physiological conditions, such as salivary flow rate, salivary composition, oxygen availability, and mechanical abrasion during mastication, may contribute to these subtle spatial microbial differences in dental plaque. However, while such spatial differences have been detected in the microbial composition of dental plaque, it is not known whether these patterns are also reflected in dental calculus. Dental calculus represents a fully mature stage of oral biofilm development that is often disrupted in living individuals practicing oral hygiene, leading to a distinct microbial profile between dental plaque and dental calculus (Velsko *et al.* 2019; Kazarina *et al.* 2021). Overall, dental calculus typically contains higher proportions of late colonizer taxa that thrive in the anaerobic environment created as the biofilm matures, and thus its composition may be less spatially variable than developing plaque biofilms, which are more dynamic and subject to periodically disruptive forces such as toothbrushing (Velsko *et al.* 2019).

However, evaluating intra-individual microbial variation in dental calculus across the dental arcade, and thus determining the degree to which a single sample can represent an individual, is challenging. Dense sampling of calculus is often hindered by missing teeth or a lack of calculus deposits distributed across the entire dental arcade. Consequently, previous studies have attempted to identify microbial spatial patterns across the dentition by instead sampling diverse individual teeth from a large number of individuals (Farrer *et al.* 2018), but this introduces a number of uncontrolled variables, such as individual differences, different biological and absolute ages of samples, different postmortem conditions, and differing degrees of preservation and degradation, which may introduce biases or otherwise alter the observable spatial patterns. Further, this approach does not allow for comparisons of how much of the variation in the dental calculus microbiome stems from intra- vs. inter-individual differences.

To determine the degree to which tooth selection matters in dental calculus sampling for comparative ancient microbiome studies, we conducted a systematic analysis of microbial spatial variation in four nearly complete human dentitions with low to heavy dental calculus deposits from the Iberian Chalcolithic site of Camino del Molino (ca. 4500-5000 BP). With dense sampling across tooth types (incisor, canine, premolar, molar) and tooth surfaces (buccal, labial, interproximal, occlusal), we performed shotgun metagenomic analysis of 87 dental calculus samples. We find that the main source of variation in the oral microbiome is the sampled individual, and therefore one randomly selected sample can, for most purposes, be used to represent an individual in population-level comparative studies. However, minor intra-individual patterns in community composition, functional potential, and species abundances are detectable with respect to tooth position (anterior vs. posterior), dental calculus deposit size, and tooth surface, although with low effect sizes. Only occlusal calculus, which is uncommon and may indicate injury or physiological dysfunction, considerably differed in composition. We found that ancient human DNA is randomly distributed across the dentition, and no spatial patterns were observed with respect to postmortem environmental contamination. Finally, we found that ancient grapevine (*Vitis vinifera*) DNA was present in the dental calculus we analyzed; however, it was also present in mandibular bone, suggesting a contaminant origin. Given that the site of Camino del Molino is located in close proximity to historic and contemporary vineyards, these findings suggest that local agricultural fields may represent a source of contamination at archaeological sites. This study contributes to an awareness of spatial variation in dental calculus microbial community composition that aims to aid researchers in developing robust study designs and valid interpretations for ancient oral microbiome studies.

## Materials and Methods

### Samples

Dental calculus was collected from four Chalcolithic (4500-5000 BP) individuals from the southeastern Iberian archaeological site of Camino del Molino near the city of Caravaca de la Cruz in Murcia, Spain, excavated during a salvage excavation in 2008 (Lomba Maurandi *et al.* 2009; Lomba Maurandi, López Martínez and Ramos Martínez 2009; Haber-Uriarte, Avilés-Fernández and Lomba-Maurandi 2011). The Camino del Molino communal burial is a natural pit with a 7 meter diameter circular base and a depth of 4 meters (of which only the lower 2 meters were used for burial), which was likely covered and sealed by a perishable structure (Lomba Maurandi *et al.* 2009). The upper layers of the site were destroyed in the early 20^th^ century as a result of agricultural terracing, but the damage did not extend to the burial deposits. Approximately 1,300 human individuals representing a broad demographic profile were buried at the site between 2800-2400 BCE (Haber-Uriarte, Avilés-Fernández and Lomba-Maurandi 2011). The site was chosen for this study because prior dental calculus research at the site had shown excellent oral microbiome preservation (Ziesemer *et al.* 2015; Mann *et al.* 2018), and microfossil studies of the dental calculus had been conducted (Power *et al.* 2014), and because the large number of individuals excavated from the site made it possible to select suitable individuals with nearly complete dentitions and sufficient dental calculus for this study. The four selected individuals were adults and had dental calculus present on most teeth (Figure 1, S1), allowing near comprehensive sampling. Dental notation below follows the FDI World Dental Federation standard (Peck and Peck 1993); molar enamel wear is reported as a Brothwell score from 1 (none) to 7 (obliteration of crown and wear of roots) (Brothwell 1972), and dental calculus deposits are graded from 1 (slight) to 4 (gross) according to Dobney and Brothwell (1987).

**Figure 1.**
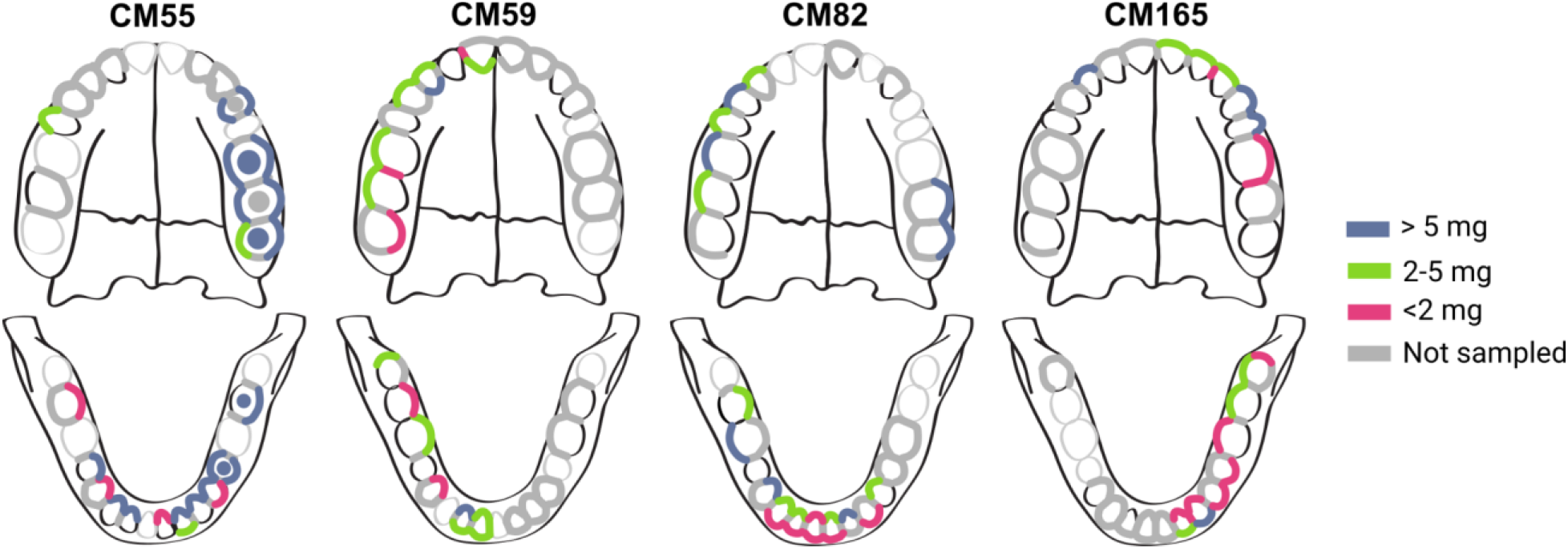
Study sampling design. Dental calculus deposits investigated in this study are highlighted and correspond to the sampled tooth surface (buccal, lingual, interproximal, occlusal). The color of the highlighting indicates the initial mass of the dental calculus deposit on the teeth that were analyzed: <2 mg (pink); 2-5.0 mg (green); >5.0 mg (blue). Teeth that were present are indicated in black outline; teeth that were absent are indicated in light gray outline. Dental calculus that was present but excluded from analysis (due to sampling design or insufficient starting mass) is marked in dark gray.

#### Individual CM55

Individual CM55 (35-39 year old female) had a complete mandible and a partial, fragmented maxilla, with a total of 22 teeth (Figure 1). Alveolar bone loss and reactive bone formation was observed throughout the mandibular periodontium, suggesting generalized periodontitis. Gross carious lesions were present in teeth 17, 35, 37, 45, and 47. Molar enamel wear was low (Brothwell stage 2). Dental calculus deposits were grade 1-2 in size, except on left premolars and molars, where they reached grade 4. The excessive calculus accumulation on the left posterior teeth, including on the occlusal surfaces, suggests that this individual had experienced pain on the left side of the mouth and avoided mastication on this side. Although no skeletal trauma was apparent, CM55 had experienced antemortem tooth loss of teeth 36 and 38, and a large carious lesion was present in 37. Significant alveolar recession and reactive bone formation was also evident around 24, but damage to the left maxilla prohibited further inspection of the bone supporting the upper molar teeth.

#### Individual CM59

Individual CM59 (25-35 year old male) had an intact mandible and a partial, fragmented maxilla, with a total of 25 teeth (Figure 1). Molar enamel wear was minimal (Brothwell stage 1-2), and no gross carious lesions were observed. Dental calculus deposits were grade 1-2 in size. Alveolar recession was slight across the periodontium, and in general the individual exhibited good dental health.

#### Individual CM82

Individual CM82 (35-45 year old female) had a complete mandible and a partial, fragmented maxilla, with a total of 23 teeth (Figure 1). Heavy enamel wear (Brothwell stage 4) was observed on the molar teeth. Dental calculus deposits were grade 1-2 in size. A large bone abscess was present adjacent to the healed alveolar bone where teeth 37 and 38 had been lost antemortem. Alveolar recession was pronounced around the molars, and healing was incomplete for four molars that had been lost antemortem. Gross carious lesions were present in teeth 16, 18, 45 and 46.

#### Individual CM165

Individual CM165 (25-30 year old likely female) had a near complete mandible and maxilla, with a total of 29 teeth (Figure 1). Although an adult, the individual retained deciduous tooth 52 and the corresponding adult tooth 12 was absent, suggesting agenesis. CM165 also had a partially impacted tooth 38. Gross carious lesions were present in teeth 37, 38, and 48. Postmortem bone breakage made the alveolar margin difficult to assess, but where observable recession was not pronounced. Molar enamel wear was low (Brothwell stage 2-3), and dental calculus deposits were grade 1 in size. Overall, the individual exhibited relatively good dental health.

Dental calculus collection was performed in an ancient DNA cleanroom environment at the University of Oklahoma (individual CM55) and the Max Planck Institute for Human History (individuals CM59, CM82 and CM165) under sterile conditions following Warinner, Velsko and Fellow Yates (2019), and supragingival dental calculus was separately collected from four different surfaces on each tooth: lingual, buccal, interproximal and occlusal. For each individual, a bone sample (approximately 50 mg) was also collected from the mandibular ramus to be used as a control for microbes characteristic of the local burial environment. As bone is mainly assumed to be free of microbes during life, in the absence of disease, the microbes identified from archaeological bone stem from the burial environment, and represent taxa that have colonized the remains, including dental calculus, postmortem. A subset of dental calculus samples was selected from each individual for metagenomic analysis (Figure 1). This subsampling was performed with the aim of achieving a balanced representation of dental sites and surfaces across individuals, as well as a consistent sample mass for analysis. For all individuals, dental sites or surfaces with < 1mg of dental calculus were generally excluded from analysis. For individuals CM59, CM82, and CM165, half of the dentition was sampled (right or left, depending on completeness and calculus abundance), but samples from the paired left/right side were also included as needed to balance out the sampling scheme with respect to tooth site and sample mass; this was particularly necessary for CM82. For individual CM55, dental calculus across the entire available dentition was sampled. Although dental calculus was mostly present only on the tooth buccal and lingual surfaces, the massive calculus deposits on the left molars of CM55 enabled the analysis of occlusal calculus for this individual. In addition, eight interproximal sites in CM59 and CM165 yielded sufficient calculus for analysis and were also sampled. In total, 87 calculus samples were selected from the four individuals for metagenomic analysis (Supplementary Data 1).

### Laboratory methods

Surface contamination was reduced by UV irradiation (30 s on both sides), followed by a washing step in 1 mL of 0.5 M EDTA (without incubation). DNA was extracted from the calculus and bone samples using a modified version of (Dabney *et al.* 2013) adapted for dental calculus (Mann *et al.* 2018; Aron *et al.* 2020) and allowing for potential future protein extraction from the same samples (Fagernäs *et al.* 2020). Briefly, the samples were decalcified in 1 mL 0.5 M EDTA for three days, after which the cell debris pellet and 100 μl of the supernatant was frozen at −20 °C and set aside for future analyses (Fagernäs *et al.* 2020). To the remaining 900 μL supernatant, proteinase K (Sigma-Aldrich) was added, and the samples were incubated at room temperature overnight. The supernatant was then mixed with binding buffer (5 M guanidine hydrochloride, 0.12 M sodium acetate, 40% isopropanol) and DNA was purified using a High Pure Viral Nucleic Acid kit (Roche Life Science) according to the manufacturer’s instructions. DNA was eluted in Qiagen EB buffer, to which Tween 20 had been added to a final concentration 0.05%. DNA was quantified using a Qubit HS assay (Thermo Fisher Scientific). Extraction blanks (one per batch) were processed alongside the samples. The full extraction protocol is available at (Aron *et al.* 2020).

Extracted DNA was processed with a partial uracil-DNA-glycosylase treatment (Rohland *et al.* 2015; Aron, Neumann and Brandt 2020) and was prepared into double-stranded libraries with dual indexing (Meyer and Kircher 2010; Kircher, Sawyer and Meyer 2012; Stahl *et al.* 2019). Library blanks were processed alongside the samples, one per batch. The DNA libraries were shotgun sequenced on an Illumina NextSeq with 75-bp paired-end chemistry. Dental calculus samples were sequenced to a depth of 10.1 ± 1.4 M reads (average ± standard deviation), bone samples to 6.6 ± 2.0 M reads, and blanks to 1.7 ± 0.7 M reads.

### Data analysis

#### Preprocessing

The EAGER v.1.92.56 (Peltzer *et al.* 2016) pipeline was used for preprocessing of the raw data. Adapter removal and merging of reads were performed using AdapterRemoval v. 2.3.1 (Schubert, Lindgreen and Orlando 2016). The reads were mapped to the human reference genome HG19 using BWA v. 0.7.12 (Li and Durbin 2009) with default settings (−l 32, −n 0.01), and unmapped reads were extracted with SAMtools v. 1.3 (Li *et al.* 2009) for downstream microbiome analyses. The unmapped reads were aligned to a custom RefSeq database (Fellows Yates *et al.* 2021) using MALT v. 0.4.0 (Herbig *et al.* 2016) (settings −id 85.0 −top 1 −supp 0.01). This database contains all bacterial and archaeal assemblies at scaffold/chromosome/complete levels (as of November 2018), with max 10 randomly selected genomes per species (prioritizing more complete genomes), as well as the human HG19 reference genome. A preliminary screening for eukaryotic DNA was also performed as described above, using the NCBI full nt database (as of October 2017), but the custom RefSeq database was chosen for further analyses, as it has been shown to yield a higher percentage aligned sequences for dental calculus (Fellows Yates *et al.* 2021). OTU tables with summarized read counts at genus level were exported through MEGAN v. 6.17.0 (Huson *et al.* 2016) (Supplementary Data 2 and 3). The R-package decontam v. 1.6.0 (Davis *et al.* 2018) was used to identify putative laboratory and environmental contaminants from OTU tables, using the prevalence method with two sets of controls (cutoff 0.8 for each): mandibular bone from the sampled individuals in this study and previously published bone samples from Bronze Age Mongolia (Jeong *et al.* 2018; Fellows Yates *et al.* 2021), and laboratory extraction and library preparation blanks.

#### Preservation assessment

A genus-level OTU table was used as input for SourceTracker v.1.0.1 (Knights *et al.* 2011). Included were also comparative samples from published shotgun microbiome studies, including 10 non-industrialized gut samples (Obregon-Tito *et al.* 2015; Rampelli *et al.* 2015), 11 industrialized gut samples (Gevers *et al.* 2012; Sankaranarayanan *et al.* 2015), 10 skin samples (Oh *et al.* 2016), 11 subgingival and 10 supragingival plaque samples (Gevers *et al.* 2012), 10 archaeological bone samples (Fellows Yates *et al.* 2021), 10 modern dental calculus samples (Fellows Yates *et al.* 2021) and 10 archaeological sediment samples (Slon *et al.* 2017). During the SourceTracker analysis, the samples were rarefied to 10,000 reads, with a training data rarefaction of 5,000. A principal component analysis was conducted on summarized genus level read counts of all samples, blanks and sources (including an additional 9 modern dental calculus samples). Multiplicative zero replacement was conducted using the R-package zCompositions v. 1.3.4 (Palarea-Albaladejo and Martín-Fernández 2015) and the data was CLR-transformed (Gloor *et al.* 2017). The non-human DNA sequences were also mapped to the *Tannerella forsythia* representative genome (strain 9212) using EAGER v. 1.92.38 as described above. The output from DamageProfiler v. 0.3.10 (Neukamm, Peltzer and Nieselt 2020) was used to visualize damage curves for the samples, and fragment length was extracted from the output table from EAGER.

#### Community composition

Analyses of community composition were conducted on the MALT taxon tables, where putative contaminants had been removed, following recommendations for compositional data (Gloor *et al.* 2017). Significant differences in community composition of samples in selected metadata groups were tested using a PERMANOVA with the R-package vegan v. 2.5.6 (Oksanen *et al.* 2019), using euclidean distance and 9999 permutations, and individuals as strata when needed. A PCA was conducted as described above. Alpha diversity was analyzed using a species-level OTU table, and Shannon Index and Inverse Simpson Index were computed using the R package microbiome v. 1.8.0 (Lahti and Shetty 2012).

#### Differential abundance

Differential abundance of species was calculated using Songbird v1.0.1 (Morton *et al.* 2019) (--formula “Jawbone+ToothSurface+ToothPosition+DepositMass_scaled+Individual”, --epochs 10000 and -- differential-prior 0.5). Tensorboard v. 1.14.0 was used for model checking. Input was a species-level OTU table, where putative contaminants were removed. Further, taxa present in fewer than three samples per individual were removed, and thereafter taxa absent in one or more of the individuals. This stringent filtering was applied in order to avoid any potential remaining contaminants or mismapping to influence the results. Two separate analyses were conducted, one without occlusal samples, and one including occlusal samples.

#### Functional analysis

The functional profiles of the microbial communities were extracted from the non-human DNA sequences using HUMAnN v. 2.8.0 (Franzosa *et al.* 2018), using the CocoPhlAn nucleotide database and the UniRef90 protein database. The output was normalized to copies per million, and translated into KEGG orthologies. Gene families were analyzed, without taking into account species assignments, and putative contaminants were removed from the dataset using decontam as described above (threshold 0.5 for both blanks and bones). A PCA was conducted, and drivers of variation identified using PERMANOVA, all as described above for community composition.

#### Human reads

In order to investigate the amount of host human DNA in the samples, while controlling for contaminating human DNA, the raw reads were mapped to the human HG19 genome as described above, with the exception of filtering for mapping quality (-q 37). Duplicates were removed using DeDup v. 0.12.2 (Peltzer *et al.* 2016), and the reads were filtered for a PMD (post-mortem damage) score of 3 using PMDtools v.0.6 (Skoglund *et al.* 2014), thereby only retaining damaged ancient reads. This is likely an underrepresentation of the number of ancient reads, as not all DNA fragments will have damage. However, assuming a consistent rate of postmortem damage accumulation over the dental arcade, the bias will be even across all sampling sites, and the patterns of damaged reads can be assumed to also represent patterns of total endogenous human reads. It was noted that occlusal samples generally have a higher percentage damage than other samples, and were therefore excluded from this analysis, as they break the assumption of equal damage. Deposit mass was accounted for in the analysis, as a positive correlation was found between deposit mass and DNA damage.

#### Plant DNA

During preliminary eukaryotic screening of the dental calculus samples, it was observed that the samples contain a considerable amount of DNA mapping to grapevine (*Vitis vinifera*). To further explore this pattern, the complete experimental dataset of dental calculus, mandibular bone controls, and blanks were mapped to the grapevine representative genome (GCA_000003745.2 12X) using EAGER as described above, with mapping quality set to 37. Damage profiles, specifically cytosine to thymine (C to T) transitions typical for ancient DNA, were created using DamageProfiler v.0.3.10 (Neukamm, Peltzer and Nieselt 2020).

#### General statistics

Unless otherwise stated, data was processed in R v. 3.6.1 (R Core Team 2019), using packages tidyverse 1.3.0 (Wickham *et al.* 2019), ggpubr v.0.3.0 (Kassambara 2018), readxl v.1.3.1 (Wickham and Bryan 2019), janitor v.2.0.1 (Firke 2018), and ggeffects v.0.14.3 (Lüdecke 2018). In order to investigate patterns across the dentition, linear mixed-effects models (LME) were fitted to the variables in question using lme4 v.1.1.23 (Bates *et al.* 2015), with the individual as the random effect when required. Model selection was performed with ANOVA using lmerTest v.3.1.2 (Kuznetsova, Brockhoff and Christensen 2017) and Box-Cox transformations identified using MASS v.7.3.51.4 (Venables and Ripley 2002). Explanatory variables in all tests are: jawbone (mandible/maxilla), tooth surface (lingual/buccal/interproximal/occlusal), tooth position (anterior/posterior), and mass of the original calculus deposit (scaled and centered continuous variable). Incisors and canines are treated as anterior teeth; premolars and molars were treated as posterior teeth. Unless otherwise noted, occlusal calculus, which was only obtained from a single individual, was excluded from most analyses because these samples were found to break the assumption of homogeneous distribution of variance (euclidean distances, ANOVA, p=0.001). 2D illustration of DNA yield, human DNA, and environmental contamination across the dental arcade was performed with ‘ili (Protsyuk *et al.* 2018), and can be accessed at https://tinyurl.com/eyjcs674. All R Markdown files have been deposited at https://github.com/ZandraFagernas/dental_arcade.

## Results

### Preservation and authentication

Total DNA yield from a sample, normalized by the mass of the dental calculus sample used for DNA extraction, may vary depending on preservation and organic matter content of the sample, and may bias downstream taxonomic profiles (Fagernäs *et al.* 2020). Occlusal samples were excluded from this analysis, as it was noted during sampling that their consistency was different from all other samples. Using linear mixed effects modeling, we tested whether tooth surface, tooth position, jawbone or deposit mass influenced the mass-normalized DNA yield from a sample. We found that none of these factors outperformed the null model (LME, individual as random effect), and therefore normalized DNA yield cannot be predicted by these variables (Figure 2A).

**Figure 2.**
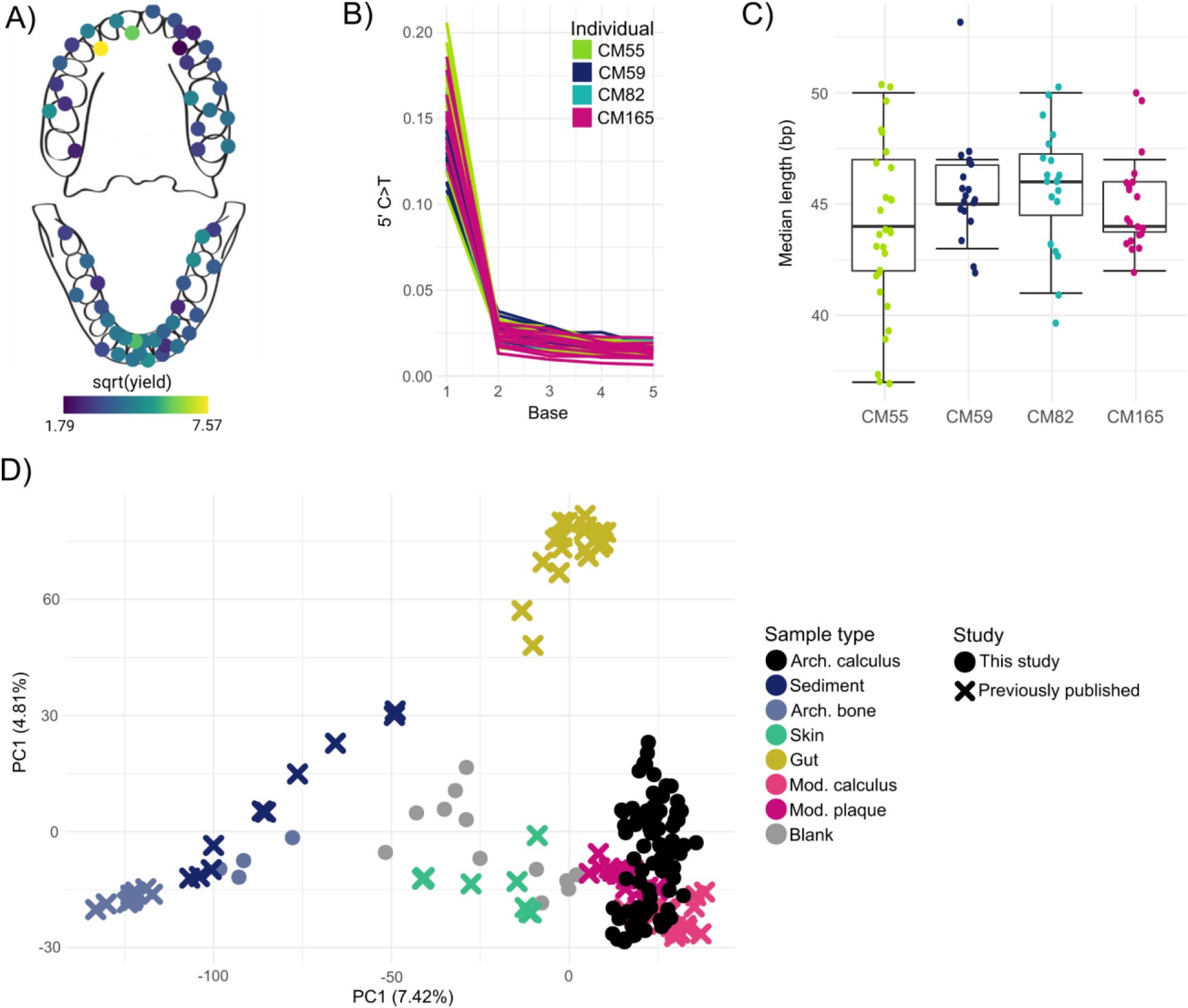
Preservation assessment of dental calculus samples. (A) normalized DNA yield (in ng DNA per mg calculus) across the dental arcade averaged across individuals. (B) C to T transitions at the 5’ end of DNA fragments aligning to *Tannerella forsythia*, consistent with ancient DNA. Note that the sharp drop from the first to the second base is due to treatment with uracil-DNA-glycosylase. (C) DNA aligning to *T. forsythia* has short median fragment lengths, consistent with ancient DNA. (D) PCA on genus level read counts of samples from this study, before removing putative contaminants; dental calculus from this study forms a cluster overlapping with modern plaque and calculus, indicating good oral microbiome preservation.

Prior to oral microbiome analysis, the archaeological dental calculus in this study was first evaluated for preservation and authenticity of the ancient oral microbiome. This is important because poor dental calculus preservation and contamination with environmental microbes can bias or interfere with downstream analyses. A PCA on genus level read counts shows that all the archaeological dental calculus samples cluster together with modern dental plaque samples, and are clearly separated from archaeological bone, gut and sediment samples (Figure 2D). To further assess preservation of the dental calculus samples, the contribution of different source environments to the composition of the samples was estimated using SourceTracker (Knights *et al.* 2011). All samples were estimated to have a majority contribution from oral microbiome sources, indicating good preservation of the oral microbiome (Figure S2). Some samples were estimated to have a minor contribution from the skin microbiome. Minor estimated contributions from the gut microbiome and sediment are also present, but are expected because gut and oral taxa are similar and can be difficult to distinguish using short read data, and because archaeological samples typically contain some soil contamination, even after washing. After taking these factors into consideration, all dental calculus samples were determined to be sufficiently well preserved for inclusion in downstream analyses.

We next assessed DNA damage patterns in the dental calculus as an indicator of authenticity. DNA from archaeological samples accumulates specific forms of damage over time, which can be seen as C to T transitions at the ends of DNA fragments and a high degree of DNA fragmentation (Dabney, Meyer and Pääbo 2013). We generated a damage plot for fragments mapping to the prevalent oral bacterium *Tannerella forsythia* (Figure 2B), and all four individuals exhibit damage patterns typical for ancient DNA that has undergone partial UDG-treatment (Rohland *et al.* 2015). The fragment length distributions of reads mapping to *T. forsythia* show that most samples have a median length <50 bp, as is expected for ancient samples (Figure 2C). Thus, taken together, the microbial DNA present within the dental calculus of the four Chalcolithic individuals in this study is consistent with an ancient and endogenous oral microbiome.

### Community composition

To determine whether local environmental and spatial variables along the dental arcade influence microbial community composition, we analyzed patterns of variation with the dental calculus samples. We found that the main driver of variation in a genus-level PCA was the individual from whom the sample originated (PERMANOVA, p<0.001, R^2^=0.14; Figure 3A), indicating that the main differences in community composition are found between individuals. When controlling for the variation introduced by the individual, the mass of the calculus deposit and tooth position (anterior *vs*. posterior) were also found to be significant drivers of variation in community composition (PERMANOVA, individual as strata, p=0.040 and R^2^=0.024 for mass; p=0.021 and R^2^=0.032 for tooth position; Figure 3B). However, although the differences are statistically significant in this study, when doing population-level comparisons they are unlikely to cause biases, as the R^2^ values are very low.

**Figure 3.**
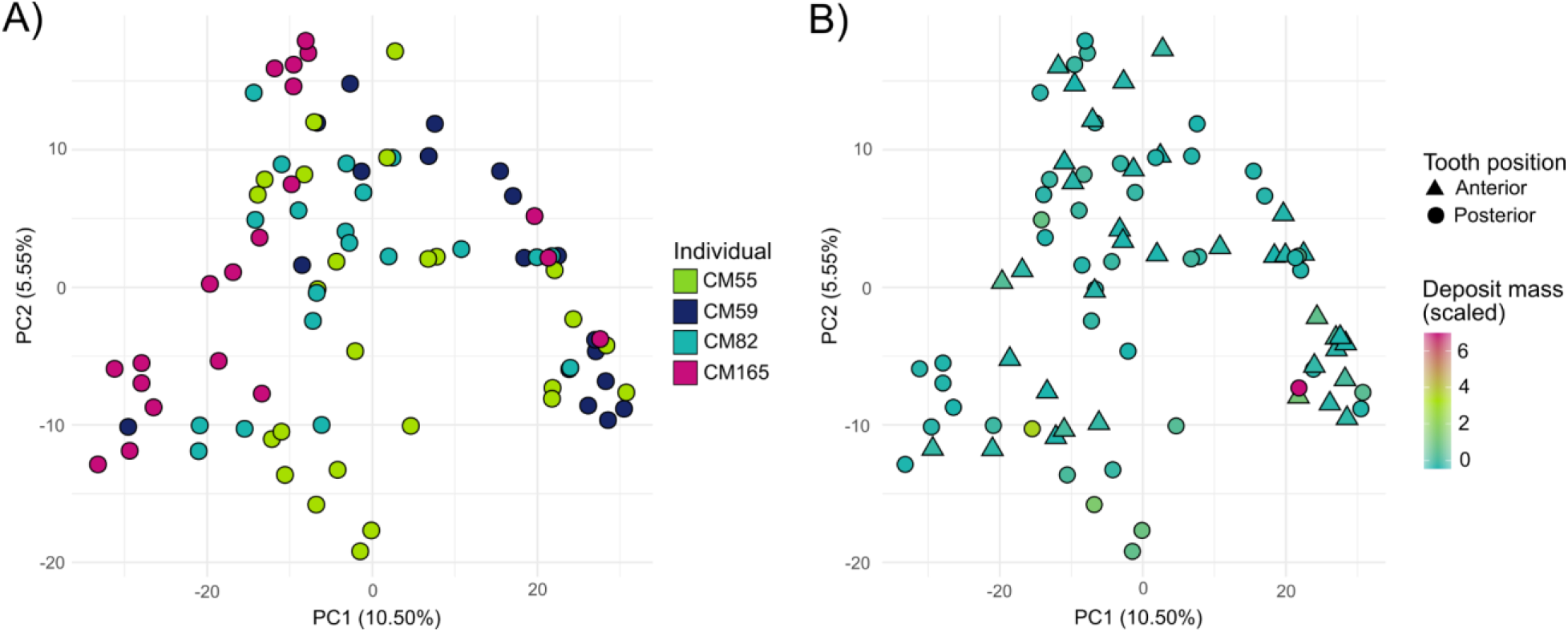
PCA on genus-level read counts. (A) All dental calculus samples plotted together, and coloured by individual. (B) Same data as A, but with samples coloured by initial deposit mass (scaled variable) and shapes representing tooth position.

We next examined alpha diversity within the dataset. Alpha diversity is a measure of how rich in species the community in a certain sample is, which may be of importance when selecting samples for a community composition study. Using the inverse Simpson Index, the mass of the original calculus deposit was found to be a significant predictor of diversity in the dental calculus samples (LME, individual as random effect, p=0.025), with diversity slightly increasing with deposit mass (Figure S3). In contrast, the null model fits the Shannon Index best, indicating that alpha diversity does not vary across the oral cavity for any of the tested variables. The Simpson index takes into account evenness, and is less influenced by rare species than the Shannon Index, indicating that the generally large number of rare species in archaeological dental calculus may erode any spatial patterns in alpha diversity.

### Differential taxonomic abundance

Due to different local environmental conditions in different areas of the oral cavity, small differences in microbial composition have been reported across the dentition in present-day dental plaque (Haffajee *et al.* 2009; Simon-Soro and Tomás 2013; Proctor *et al.* 2018). It is, however, not known if such patterns can be detected in archaeological samples, after both biofilm maturation during life and postmortem degradation over time. Here, we find that there are slight taxonomic differences with respect to tooth surface and initial deposit mass. First, we observe differences in taxa between anterior (incisors and canines) and posterior (premolar and molars) teeth, where the taxa that are more abundant in the anterior teeth are more often aerobic or facultatively anaerobic, while the taxa that are most associated with posterior teeth are anaerobic (Figure 4A). Second, interproximal spaces seem to be enriched in species belonging to the genera *Methanobrevibacter* and *Olsenella* (Figure 4B), which are both acid tolerant anaerobes. Finally, the species *Actinobaculum* sp. oral taxon 183 and *Fusobacterium* sp. oral taxon 203 are found at a higher abundance in low mass dental calculus deposits, as compared to high mass deposits (Figure 4C). However, little is known about the physiology or role in the dental plaque biofilm of these taxa. Fusobacteria are generally secondary colonizers in the dental plaque biofilm, and bind to several other bacterial taxa (Kolenbrander 1988). A 2D model showing the spatial distributions of the taxa in Figure 4A–C across the dentition can be found at https://tinyurl.com/eyjcs674.

**Figure 4.**
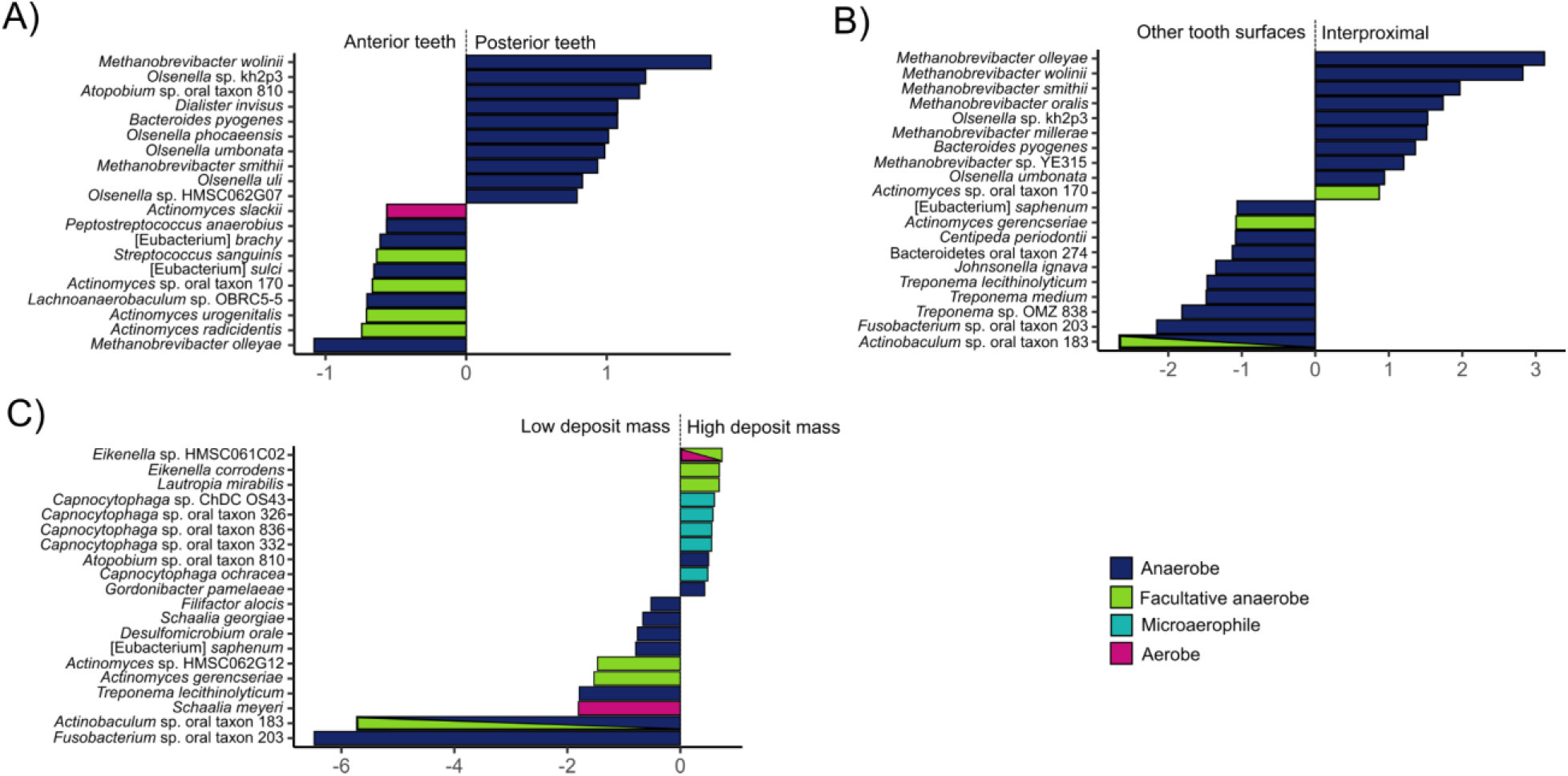
Differential abundance of species across the oral cavity. A) Species associated with posterior (premolars and molars) vs. anterior (incisors and canines) teeth. B) Species associated with interproximal spaces vs. all other tooth sites. C) Species associated with high vs. low initial deposit mass. Only the top ten taxa most associated with each factor are shown.

### Functional profile

In addition to their taxonomic composition, microbial communities may also differ in their gene content, and therefore functional potential. It has been seen that although microbial community composition may be similar between individuals, they can differ in the functional potential of the microbiome (Fellows Yates *et al.* 2021). To evaluate whether there are potential functional differences across the dental arcade, we analyzed the genes present in the dental calculus metagenomes. In total, 2,791 gene families were identified in the dataset, after removing putative contaminants that were identified from blanks and bone samples. The individual was found to be the strongest driver of variation (PERMANOVA, p=0.001 and R^2^=0.12; Figure 5), and after accounting for this, tooth surface (p=0.020, R^2^=0.049), tooth position (p=0.039, R^2^=0.029), and deposit mass (p=0.013, R^2^=0.036) were found to significantly drive functional variation (PERMANOVA, individual as strata). However, these factors explain only a very minor part of the variation, as can be seen by the low R^2^ values. It should also be noted that the tooth surface variable breaks the assumption of homogeneity of variance for this analysis, which may affect the results of the PERMANOVA.

**Figure 5.**
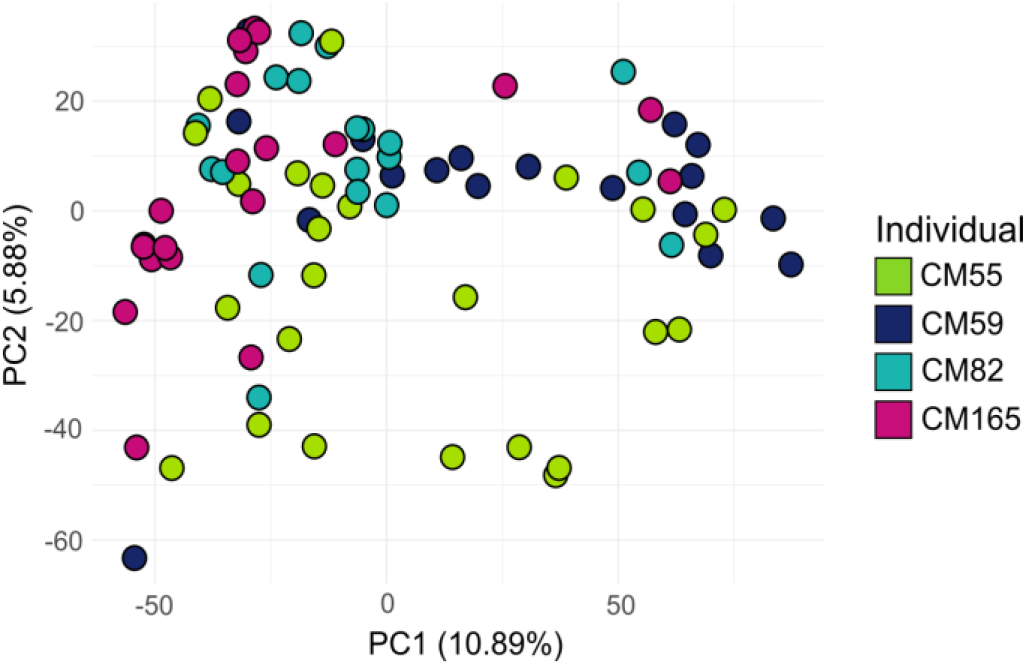
Functional profile of dental calculus samples. PCA of gene families, normalized to copies per million, with colour indicating individual.

### Human genetic content

Although dental calculus generally contains a very low proportion of human DNA (Mann *et al.* 2018), different enrichment approaches have been used to increase the human DNA fraction enough to study the human genome (Ozga *et al.* 2016; Ziesemer *et al.* 2019). Human DNA from dental calculus is mainly derived from a single individual, the host (Ozga *et al.* 2016). Human DNA may in theory be differentially incorporated into dental calculus across the dental arcade, depending on salivary flow, inflammation, or disease, among other factors. We investigated the presence and relative abundance of ancient human DNA in our samples to assess potential spatial patterning of human host DNA in calculus. To focus our analysis on host ancient DNA, we restricted our analysis to only DNA fragments with C to T DNA damage. As a slight positive correlation was found between deposit mass and damage (Figure S4), deposit mass was accounted for in this analysis. We found that the best fitting model for predicting the proportion of human reads in the dental calculus samples is a null model, indicating that the distribution of human DNA in dental calculus does not significantly vary according to tooth surface, tooth position, or jawbone (LME, deposit mass as random effect; Figure 6A).

**Figure 6.**
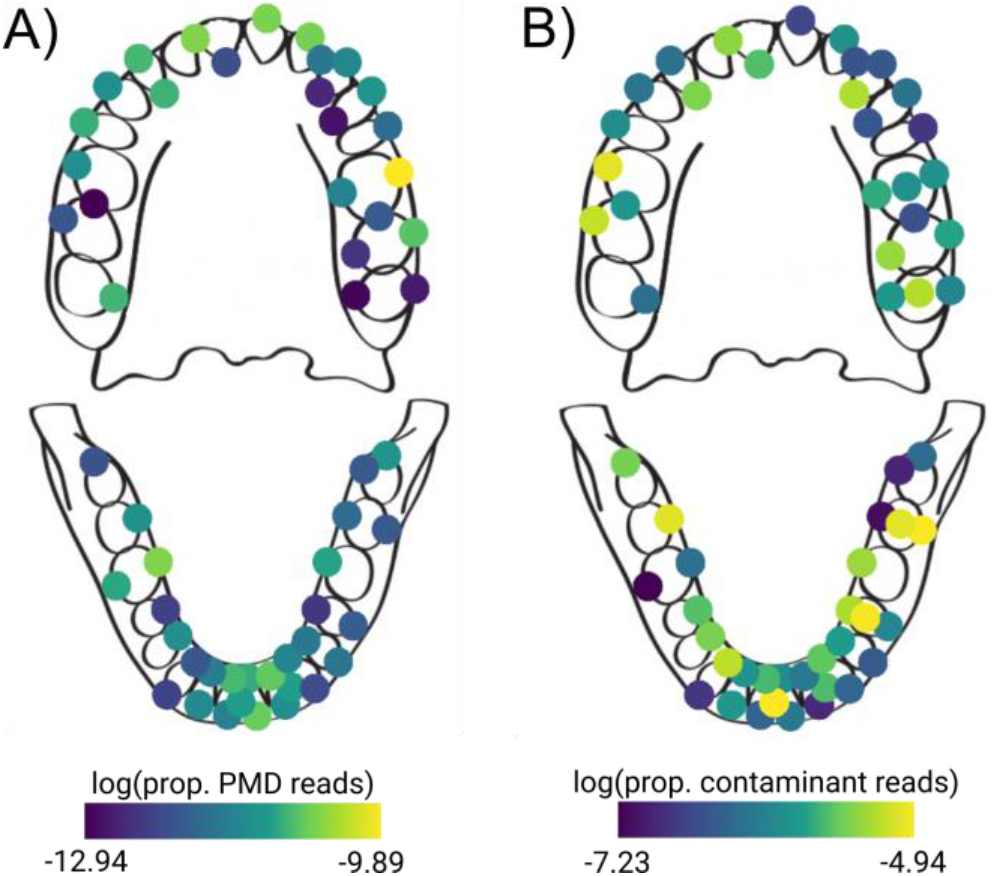
Distribution of ancient human reads and environmental contaminant reads across the dental arcade. A) Proportion of human reads with post-mortem damage (PMD) of the total number of sequenced reads, averaged across individuals for each sampling location. B) Proportion of reads that stem from putative environmental contaminant taxa out of all reads, averaged across all four individuals for each sampling location.

### Postmortem environmental colonization

Whether contamination by infiltration of environmental microbes from the burial context is introduced in a non-random way across the oral cavity is not known. Because different properties of calculus across the dentition could make certain regions more susceptible to external colonization, we investigated the distribution of environmental contaminant reads in our samples. On species level, a total of 215 taxa (out of 556 taxa) were identified as putative environmental contaminants in the entire dataset, using the bone samples from the mandibles as a proxy for colonizing microbes from the burial ground. This analysis was performed at the species level, as it is possible for taxa in the same genus to grow in different habitats. We tested whether the distribution of these species across the dentition was influenced by tooth surface, tooth position, jawbone or deposit mass using linear mixed effects modeling; however, we found that none of these factors outperformed the null model. Therefore, it appears that contamination is randomly distributed across the dental arcade (LME, individual as random effect; Figure 6B).

### Occlusal calculus

The occlusal dental calculus analyzed in this study differed from the calculus from other tooth surfaces in several ways, and was therefore excluded from most analyses. During sampling, occlusal calculus was found to have a different consistency from the other calculus, being less dense and having less structural integrity. Occlusal calculus was found to have a higher amount of DNA damage than other calculus. For reads mapping to *Tannerella forsythia*, a model including tooth surface and deposit mass best predicted damage at the 1st base at the 5’ end of the fragment (LME, individual as random effect, p=0.018), with occlusal samples having higher levels of damage than other samples (Figure S4). Further, occlusal calculus samples broke the assumption of homogeneity of dispersion for the community composition, which may be due to the fact that they were only collected from a single individual, and from only posterior teeth on the same side of the mouth. Overall, we found that despite forming on posterior teeth, occlusal calculus samples are somewhat enriched in aerotolerant species, possibly due to their more exposed location on the tooth, compared to the lingual and labial surfaces of the posterior teeth that directly abut the tongue and buccal mucosa, respectively (Figure 8).

### Plant DNA

Ancient dental calculus is a potentially valuable source of information about ancient diets, as it is possible to directly study diet-related biomolecules and microfossils incorporated in the calculus during an individual’s lifetime. Researchers have previously attempted to identify dietary sources using DNA from dental calculus (Warinner *et al.* 2014; Weyrich *et al.* 2017), an approach that also has many difficulties due to the exceptionally low number of dietary DNA sequences typically found in dental calculus (Mann *et al.* 2020). The dental calculus samples in this study contained trace amounts of plant DNA fragments (between 170-1578 reads per individual, or 0.002-0.011% of total reads) mapping to the grapevine (*Vitis vinifera*) genome, which is currently and historically widely cultivated in the region. However, it was noticed that similar numbers of grapevine reads (205-2119 reads, or 0.003-0.034%) were also recovered in the mandibular bone control samples (Figure 7A). Both sets of reads were found to have C to T damage typical of ancient DNA (7-9% for bones and 3-14% for calculus; Figure 7B), but at lower levels than observed for the oral bacterium *T. forsythia* (11-21%; Figure 2B). The presence of grapevine reads in both dental calculus and bone, together with the lower amount of damage, suggests a likely postmortem origin of the grapevine DNA. However, a dietary origin of the grapevine DNA cannot completely be excluded, as a wild variety has been documented in the region since the Palaeolithic (Aura *et al.* 2005; Iriarte-Chiapusso *et al.* 2017).

**Figure 7.**
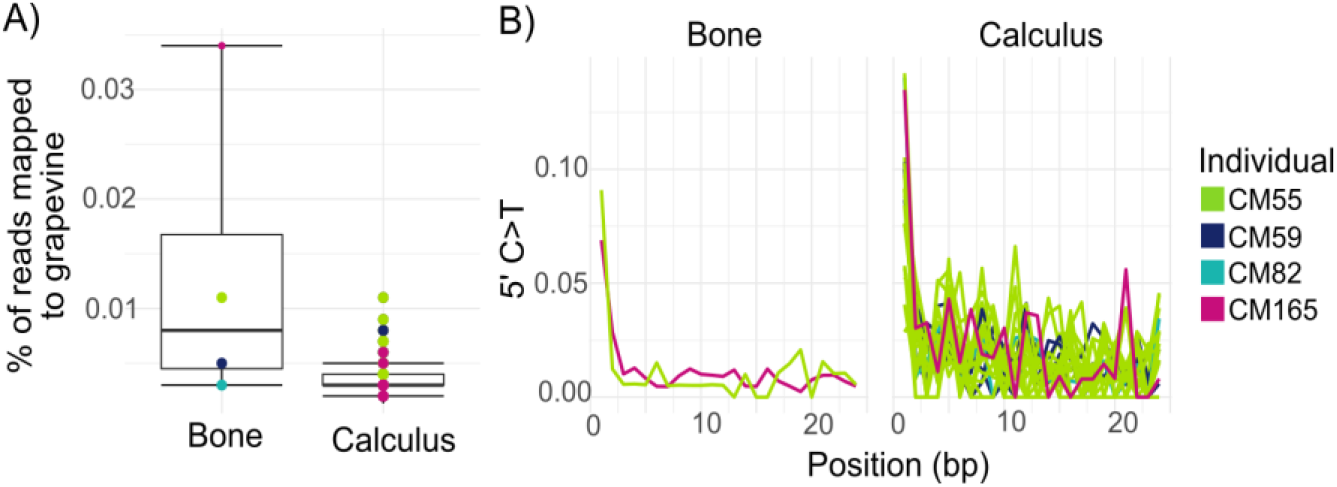
Presence of grapevine DNA in bones and dental calculus samples. A) The percentage of reads that aligned to the grapevine genome per sample. B) C to T miscoding lesions at the 5’ end of the read, for each sample with >500 reads aligning to grapevine.

**Figure 8.**
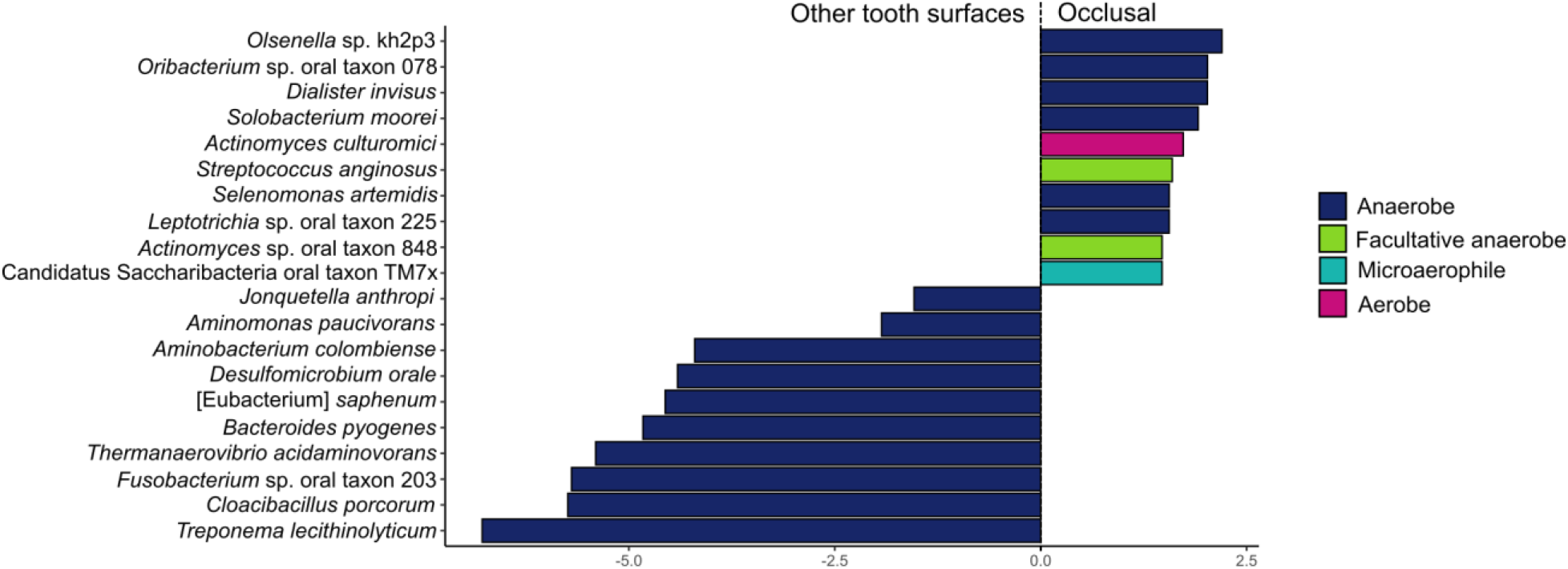
Differential abundance of taxa in occlusal samples compared to others. Only the top ten taxa are shown, and the bars are coloured by aerotolerance of the taxa.

## Discussion

A potentially uneven distribution of microbes in microbiomes can cause biases in downstream analyses if spatial variation is not taken into account during sampling design and data interpretation. Archaeological dental calculus provides a valuable window into the evolution of the oral microbiome, but to date it has not been clear to what degree microbial taxa are spatially patterned across the dentition and, thus, to what degree sampling strategy might impact comparative studies of dental calculus microbial communities. The results of present-day dental plaque studies cannot be directly applied to dental calculus because the two substrates reflect different levels of biofilm maturity and have slightly different composition (Velsko *et al.* 2019), and in previous studies of spatial variation in archaeological dental calculus, which sampled diverse individual teeth from a large number of individuals (Farrer *et al.* 2018), potentially confounding factors such as individual, temporal, environmental, and taphonomic differences were not controlled for. Here, we have presented a systematic study of intra-individual variation in archaeological dental calculus by focusing on intensive, comprehensive sampling of the dentitions of four contemporaneous individuals from the same burial context.

Overall, we find that although there are small differences in the spatial distribution of anaerobic and aerotolerant taxa, as well as minor associations between taxonomic composition and initial calculus deposit size, these factors account for very little of the overall microbial and functional variation within dental calculus. Spatial patterns in the oral microbiome that have been identified in studies of modern dental plaque (Haffajee *et al.* 2009; Simon-Soro and Tomás 2013; Proctor *et al.* 2018) are not obvious in this study. Such patterns may have been present during life but were subsequently lost over time due to taphonomy, or these patterns may simply not be present in calculus. Although taphonomic processes, such as C to T damage accumulation and DNA fragmentation, as well as postmortem colonization of the body by environmental taxa, may obscure oral microbiome spatial patterns, we did not find these factors to correlate with the microbial patterns we observed. A study of modern dental calculus that investigates species spatial patterning will be needed to determine if the patterns observed in dental plaque are maintained as the biofilm matures and calcifies into dental calculus.

Although this study investigated a small number of individuals from a single archaeological site, the purpose of this study design was to limit the number of potentially confounding factors, such as different sample ages, different burial conditions, and different storage and handling practices after excavation. Microbial spatial patterning may differ in other populations or at other archaeological sites, and this warrants further investigation. However, as the species profiles of human dental calculus appear to be more consistent across time, space, and health status than dental plaque (Velsko *et al.* 2019; Fellows Yates *et al.* 2021), it is possible that any variation will be very minor.

Although we observed few spatial patterns in archaeological dental calculus, we find that occlusal calculus may represent a special exception. Dental calculus rarely accumulates on the occlusal surfaces of teeth, in part due to the abrasive forces of mastication, and large deposits of occlusal calculus are generally indicative of physiological injury or dysfunction. Here, only one individual had occlusal calculus, but this calculus had a distinct texture, higher DNA damage, and different levels of taxonomic dispersion compared to other dental calculus in the study, even from the same individual. Although further research on a larger number of individuals is necessary, occlusal calculus is likely not representative of oral microbiome communities, and therefore should be avoided in comparative studies of microbial variation across individuals.

Beyond microbes, dental calculus is also valuable because it entraps dietary and other environmental debris during life, and thus can provide clues about the foods and activities of past societies (Hardy *et al.* 2009; Leonard *et al.* 2015; Power *et al.* 2015; Radini *et al.* 2017). Although dietary proteins have been shown to preserve within dental calculus (Hendy *et al.* 2018; Wilkin *et al.* 2020; Scott *et al.* 2021), the metagenomic recovery of dietary DNA from calculus has yielded more equivocal results (Mann *et al.* 2020). The recovery and authentication of eukaryotic DNA in metagenomic datasets is not trivial due to complicating factors such as the very low number of non-host eukaryotic DNA fragments typically found in dental calculus and the problem of microbial contamination in eukaryotic reference genomes, which can lead to false positives (Mann *et al.* 2020). Here, we show that an additional complicating factor may be contamination from the environment, and specifically from nearby agricultural fields. It is therefore advisable to include environmental controls, such as bone or sediment samples, in metagenomic studies of diet. Another authentication aid may be the use of complementary dietary identification methods, such as microfossil analysis or palaeoproteomics. Through proteomic analyses, for example, it is possible to deduce the part of an organism from which the biomolecules originate, such as seed proteins from plant seeds, or milk proteins from dairy products. Combining methods may thus aid researchers in establishing the plausibility of a given organism being incorporated into dental calculus as a food as opposed to environmental contamination.

To conclude, we find that in most applications a single sample of archaeological dental calculus can be used to represent an individual in comparative studies of the ancient oral microbiome. The use of a single sample instead of multiple samples, either pooled or studied separately, reduces the destructive demands on this finite archaeological material. However, as there are minor spatial patterns present, care should be taken to record the sampling location within the oral cavity for each dental calculus sample, whenever possible. This makes it possible to later reevaluate findings if systematic biases are suspected.

## Accession numbers

Genetic data have been deposited in the ENA under the accession PRJEB46022.

## Acknowledgements

This work was supported by the Werner Siemens Stiftung (“Paleobiotechnology” to C.W.), the European Research Council (STG–677576 “HARVEST” to A.G.H.), the Generalitat Valenciana (GV/2015/060 and CIDEGENT/2019/061 to DCSG), the Spanish government (PID2019-111207GA-I00 and EUR2020-112213 to DCSG), and the Max Planck Society. The authors thank James Fellows Yates for assistance with data analysis.

## Supplementary figures

**S1.**
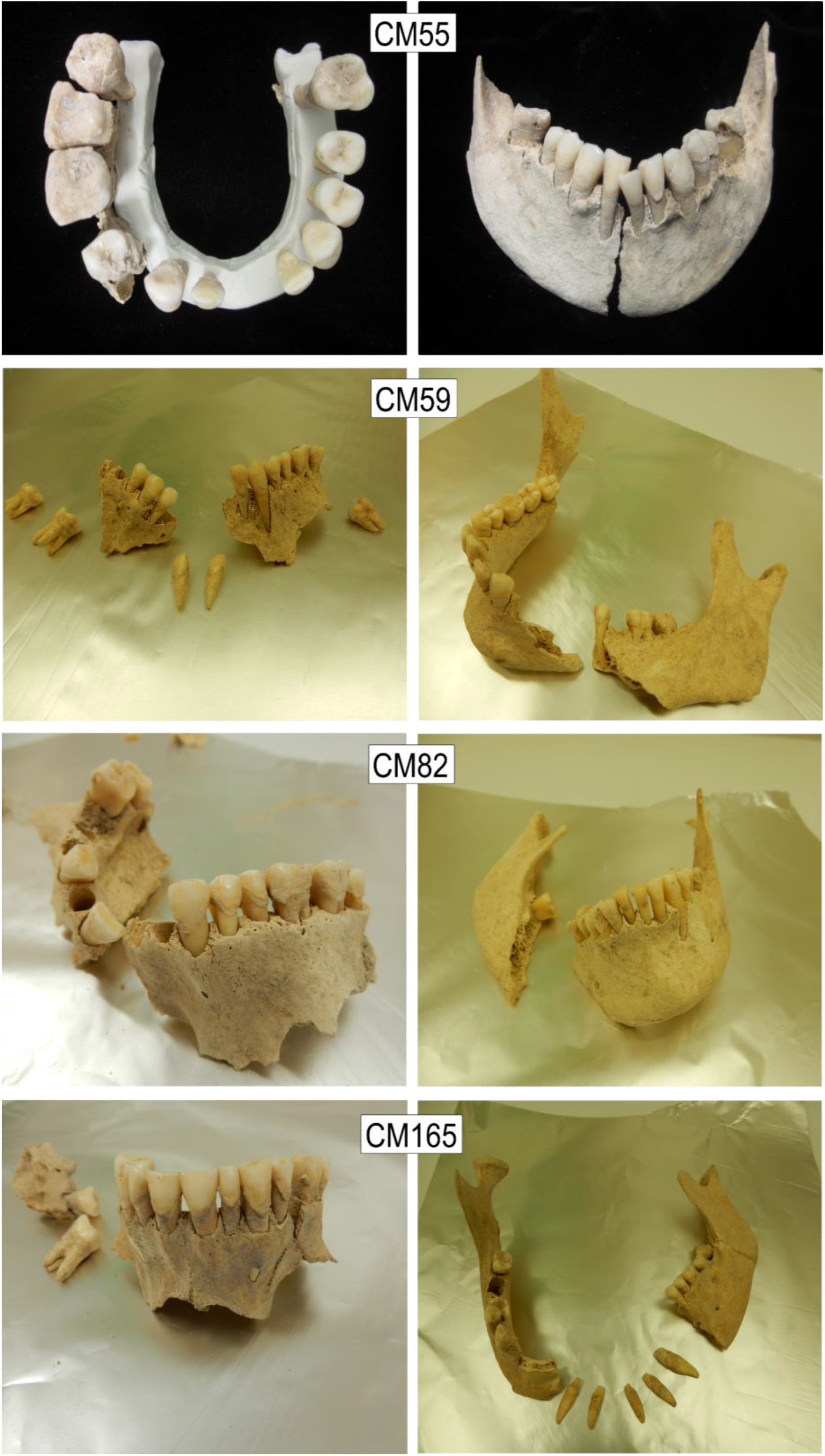
Photos of the entire available dentitions of the four individuals sampled in this study.

**S2.**
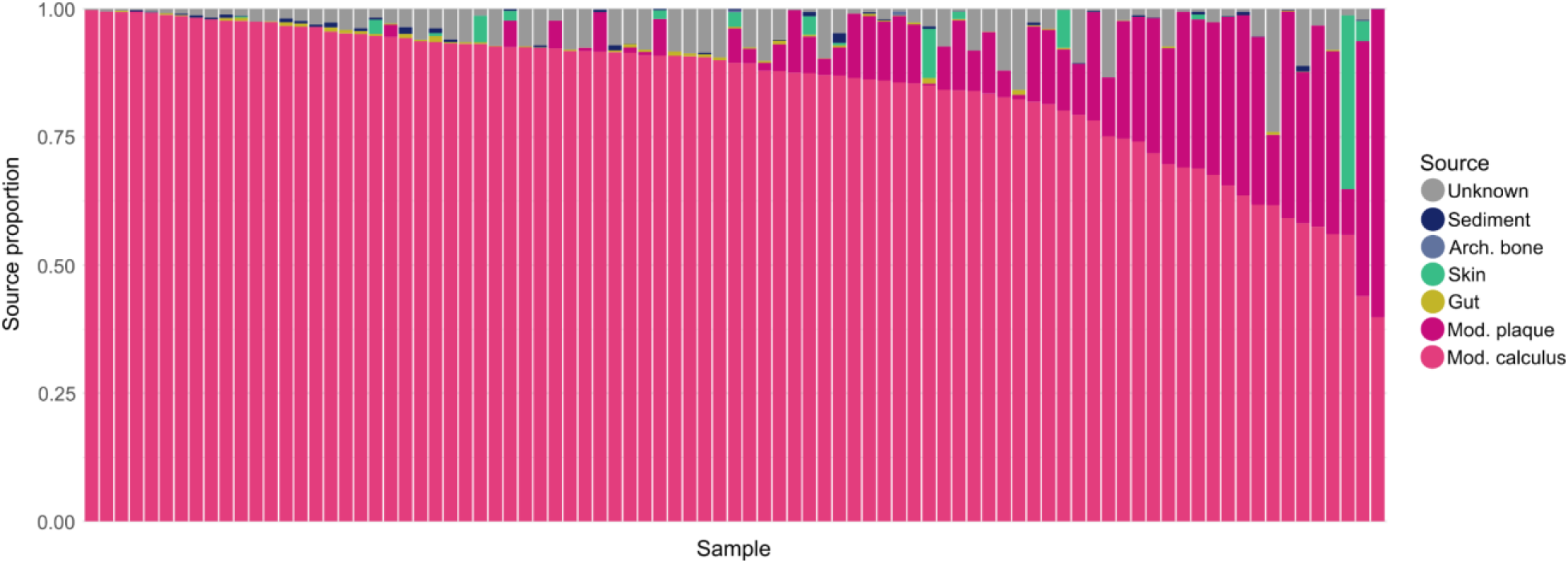
Sourcetracker results, generated from a genus-level OTU-table.

**S3.**
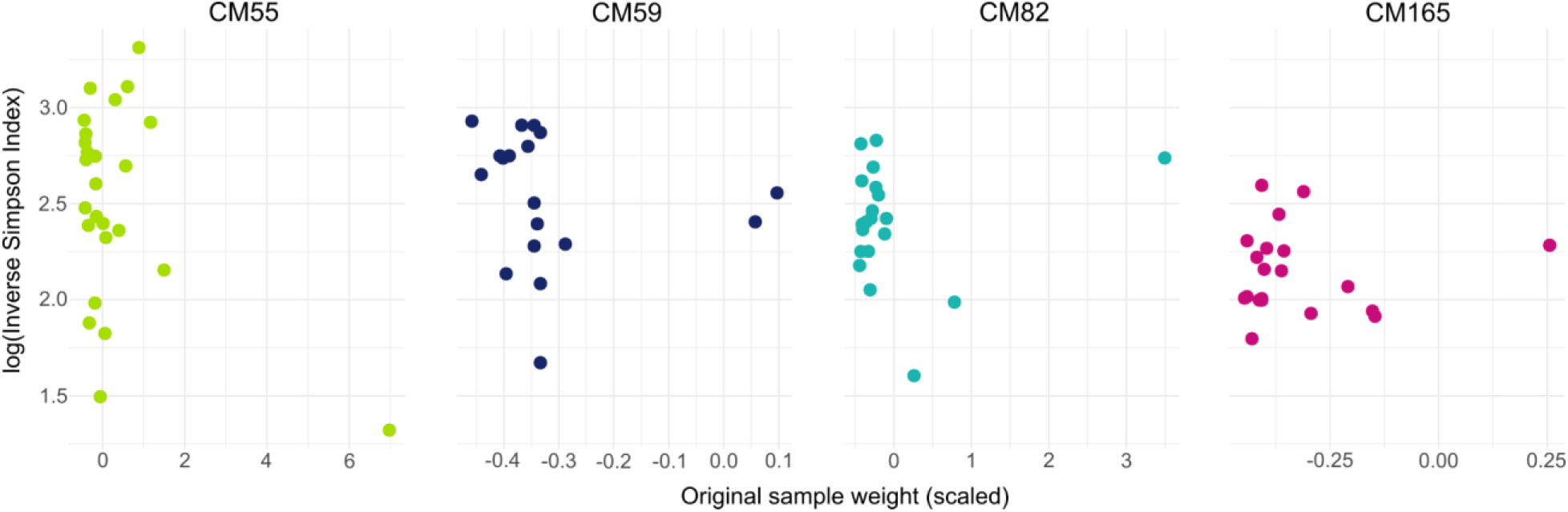
Inverse Simpson Index by mass of the original calculus deposit.

**S4.**
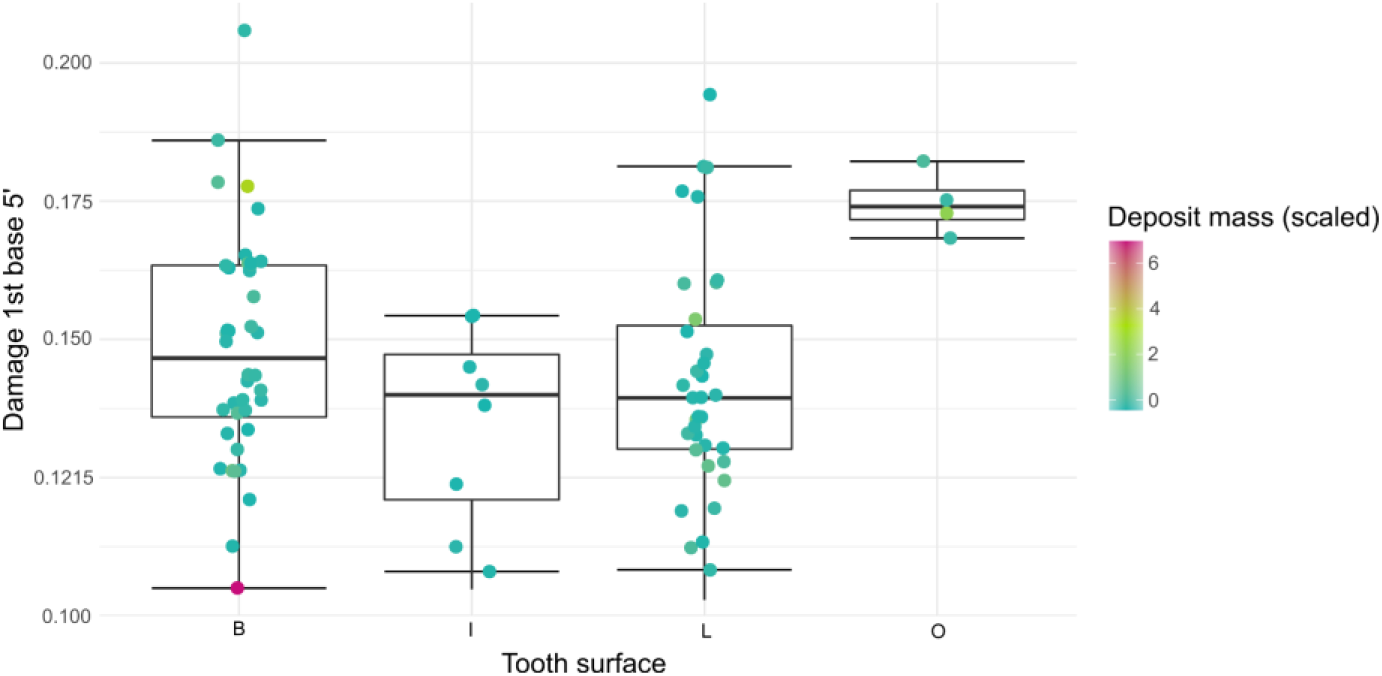
Damage of first base at the 5’ end of fragments mapping to *Tannerella forsythia*.

